# The runaway evolution of SARS-CoV-2 leading to the highly evolved Delta strain

**DOI:** 10.1101/2021.12.30.474592

**Authors:** Yongsen Ruan, Mei Hou, Xiaolu Tang, Xionglei He, Xuemei Lu, Jian Lu, Chung-I Wu, Haijun Wen

## Abstract

In new epidemics after the host shift, the pathogens may experience accelerated evolution driven by novel selective pressures. When the accelerated evolution enters a positive feedback loop with the expanding epidemics, the pathogen’s runaway evolution may be triggered. To test this possibility in COVID-19, we analyze the extensive databases and identify 5 major waves of strains, one replacing the previous one in 2020 – 2021. The mutations differ entirely between waves and the number of mutations continues to increase, from 3-4 to 21-31. The latest wave is the Delta strain which accrues 31 new mutations to become highly prevalent. Interestingly, these new mutations in Delta strain emerge in multiple stages with each stage driven by 6 – 12 coding mutations that form a fitness group. In short, the evolution of SARS-CoV-2 from the oldest to the youngest wave, and from the earlier to the later stages of the Delta wave, is a process of acceleration with more and more mutations. The global increase in the viral population size (*M*(*t*), at time *t*) and the mutation accumulation (*R*(*t*)) may have indeed triggered the runaway evolution in late 2020, leading to the highly evolved Alpha and then Delta strain. To suppress the pandemic, it is crucial to break the positive feedback loop between *M*(*t*) and *R*(*t*), neither of which has yet to be effectively dampened by late 2021. New waves beyond Delta, hence, should not be surprising.

## Introduction

Pathogens that made the jump from animal hosts may immediately experience novel selective pressures and rapid evolution in the human hosts(Parrish, et al. 2008; Plowright, et al. 2017; Cui, et al. 2019; Andersen, et al. 2020; Lytras, et al. 2021; Ruan, et al. 2021; Wu, et al. 2021). We now test the hypothesis that SARS-CoV-2 has experienced accelerated adaptive evolution in 2020 – 2021. The extensive genomic sequences of SARS-CoV-2 afford evolutionists an unprecedented opportunity to track the evolution in ways unimaginable in the study of any other living organisms. In particular, the data collection covers in real time the entire span of evolution across most geographical regions.

In addition to the host shift, there may be additional forces that could drive the accelerated evolution of SARS-CoV-2. First, the evolution of herd immunity may elicit an arms race between host and pathogen(Peschel and Sahl 2006; Guo, et al. 2020; Kajan, et al. 2020; Ruan, et al. 2021). Second, new strains may evolve and compete in the infection of human hosts(Choi, et al. 2020; Sashittal, et al. 2020; Kemp, et al. 2021). Third, with “mutations-begetting-mutations”(Ruan, Wang, Chen, et al. 2020), the mutation rate may increase dramatically, as documented in cancers. Fourth, viral adaptive evolution and viral population size may mutually reinforce each other. There is some evidence for each of these forces. Importantly, in each case, there is a positive feedback loop that would lead to the escalation of the rate of adaptive evolution. We will refer to the escalated adaptive evolution in such a loop “runaway evolution”. In particular, the feedback loop between the viral population size and the rate of adaptive evolution may be most easily tracked.

In analyzing the very large number (> 3 millions) of sequenced genomes of modest size (~ 30 Kb for SARS-CoV-2), we take two complementary approaches, referred to as the infinite-allele and infinite-site model, respectively(Kimura and Crow 1964; Kimura 1969). In the former, one treats each sequence as an allele (i.e., haplotype) and compare the alleles by, for example, constructing their genealogical relationships(Forster, et al. 2020; Rambaut, et al. 2020; Tang, et al. 2020a; Kumar, et al. 2021; Tang, et al. 2021). This infinite-allele approach, commonly used in studying viral evolution, will be used here in a preliminary probe. For the in depth analyses, we will use the infinite-site model whereby one examines each variable site across all sequences. The two models, equivalent in dynamics, reveal different aspects of the same evolutionary phenomena. Using the combined approaches, we aim to determine if SARS-CoV-2 has been experiencing accelerated adaptive evolution via the runaway process.

## Results

In Part I, we outline the hypothesis of runaway viral evolution in a positive feedback loop. In the main Part II, the genomic data of SARS-CoV-2 are analyzed to test the hypothesis.

### I. The hypothesis of runaway viral evolution

As stated in the Introduction, there are multiple factors that can accelerate the adaptive evolution of SARS-CoV-2. One of the factors that can be most easily formulated is the viral population size at time *t, M*(*t*). Other factors may be no less important but the *M*(*t*) data are readily available. We shall follow the convention of using only one prevalent strain to represent the virions in each infected host. Hence, *M*(*t*) is assumed to be equivalent to the number of infections at that time. According to the standard theory(Crow and Kimura 1970; Eyre-Walker 2006; Ruan, Wang, Zhang, et al. 2020), the rate of adaptive evolution can be expressed as

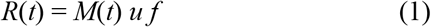

where *u* is the mutation rate and *f* is the fate of the mutation (expressed in probability). The rate of adaptive evolution, *R*(*t*), is the number of advantageous mutations produced at time *t* that will become fixed, or at least become prevalent, in the population.

If all mutations are equal in fitness, *f* = 1/*M*(*t*). Thus, the rate of neutral evolution would be *R*(*t*) = *u* and the rate is independent of the population size. For adaptive mutations, *f* is a function of the selective advantage that is often independent of *M*(*t*)(Crow and Kimura 1970; Eyre-Walker 2006; Ruan, Wang, Zhang, et al. 2020). Interestingly, a higher *R*(*t*) means more advantageous mutations that promote infections and increase *M*(*t*). Therefore, *R*(*t*) and *M*(*t*) would form a positive feedback loop as indicated by arrows of acceleration:

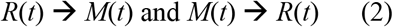

When such feedbacks are in operation, *M*(*t*) would grow like a snowball, leading to out-of-control epidemics. *M*(*t*) → *R*(*t*) should not be in dispute but the other half, i.e., *R*(*t*) → *M*(*t*), may not be true for most species. In general, adaptive evolution is not manifested as large population sizes, which are usually environmentally limited. In viral evolution, however, an increase in *R*(*t*) may be directly translated into a larger population size, given that viral populations can rapidly expand or contract by orders of magnitude.

In this backdrop, we wish to assess the possibility of COVID-19 experiencing accelerated evolution in 2021. While SARS-CoV-2 seemed to evolve slowly in the early part of 2020, there have been several waves of SARS-CoV-2 strains, including the latest Delta strain. We shall determine the genomic bases of the strains driving these waves.

### II. The analyses of SARS-CoV-2 evolution

In the sub-section 1 below, we examine the evolution of SARS-CoV-2 haplotypes, one sequence a time across sites. In the remaining subsections, we analyze the frequencies of mutants one site at a time across sequences.

#### 1. The accelerated evolution of SARS-CoV-2 alleles

Following Eq. (1), we compare the *R*(*t*) and *M*(*t*) values from the beginning of the COVID-19, set at Dec. 31, 2019 (day 0) as shown in Fig. 1. For *R*(*t*), we use the rate of non-synonymous evolution, calibrated by the synonymous changes, as the proxy. The graphs present cumulative numbers; hence the focus should be on the rate of increase, i.e., the slope of the curve.

**Fig. 1.**
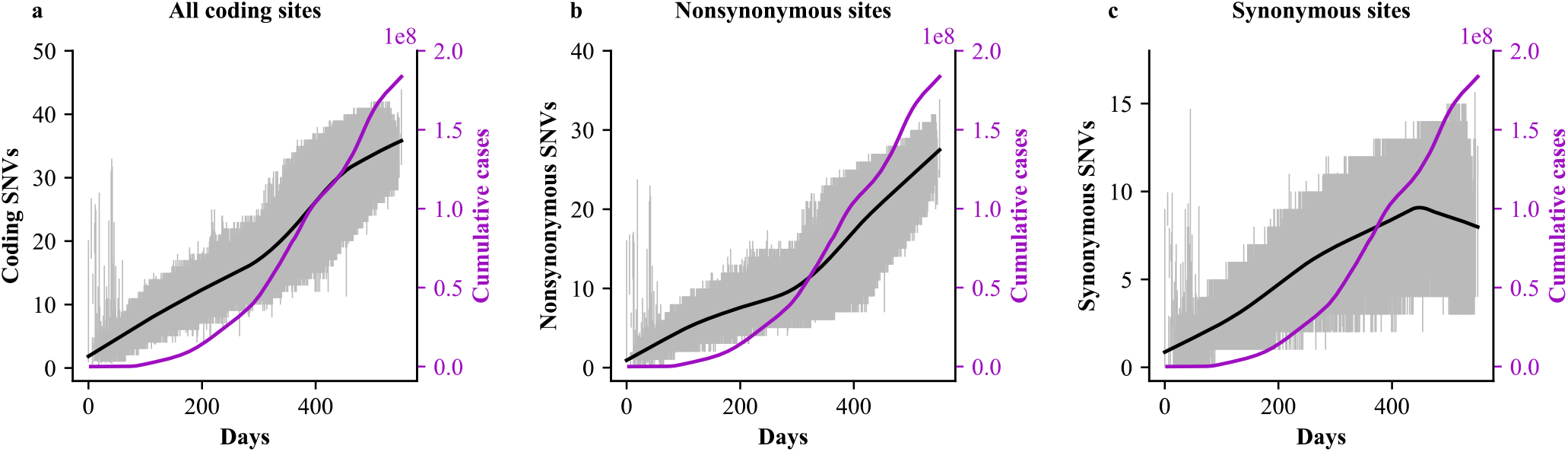
The number of SNVs accrued in the genomes of SARS-CoV-2 in the last 550 days. The number of SNVs (relative to the reference genome, y-axis) of each strain is plotted against the number of days between the collection time and the start of the epidemics (x-axis). The SNVs are, respectively, from the coding regions (**a**), non-synonymous sites (**b**) and synonymous sites (**c**). For each date, hundreds or thousands of strains were collected with the average (the black line) and 95% quantiles (the grey shade) given. The purple line is the cumulative number of confirmed COVID-19 cases worldwide (downloaded from the WHO website).

We downloaded 1,853,355 SARS-CoV-2 genome sequences with reliable information on the collection time from GISAID (https://www.gisaid.org, as of July 5, 2021). Among them, 97.8% belongs to the L lineage(Tang, et al. 2020b; Tang, et al. 2021). Hence, almost all the comparisons are between the L strains and the reference genome (Wuhan-Hu-1, GenBank: NC_045512), also of the L-type. In Fig 1a, we plotted the number of SNVs in the coding regions of each strain vis-a-vis the reference genome (*y*-axis) according to the isolation date of that strain (*x*-axis).

When one examines the total number of accrued mutations, the increase looks linear with time whereas *M*(*t*) increases sharply toward the end of 2020 (Fig. 1a). The average line is shown and the grey zone represents 95% quantiles. The patterns are more informative when the nonsynonymous and synonymous changes are separated. Fig. 1b shows that the rate of nonsynonymous changes has started to increase by late 2020. The trend appears to coincide with the number of infections, *M*(*t*), which also starts to increase at about the same time. Note that the coincidence is not observed for synonymous changes, as shown in Fig. 1c. The trend of Fig. 1b suggests that COVID-19 may have entered the runaway phase of evolution in late 2020 to early 2021 when *M*(*t*) started to accelerate. It is possible that the flu pandemic of 1918 – 1920 may have entered such a phase as well. Since COVID-19 is the first pandemic that has been tracked by large-scale viral sequencing, the runaway phenomenon can be more clearly revealed.

#### 2. The five waves of SARS-CoV-2 evolution

To study the rise of new strains, we track variant frequencies site-by-site since such sites can often be connected to functionality(Hou, et al. 2020; Korber, et al. 2020; Li, et al. 2020; Dejnirattisai, et al. 2021; Deng, et al. 2021; Planas, et al. 2021; Plante, et al. 2021; Tao, et al. 2021; Volz, et al. 2021; Zhang, et al. 2021). Furthermore, by using the site-model, one only needs to track < 100 variants sites, instead of more than one million sequences.

Fig. 2 tracks the variants that have reached 0.3 or above in frequency at its peak. In the UK samples (Supplementary Table 1), 51 nonsynonymous,16 synonymous, and 5 non-coding variants meet the criteria with an A:S:NC ratio of 51:16:5 (A for amino-acid changes, S for synonymous ones, and NC for non-coding ones). These variants emerged in five waves in the UK data, labelled W0 to W4 in Fig. 2a. The patterns from the four geographical regions (see Supplementary Table 1-4) are comparable but the waves appear more sharply defined in UK. The mutations associated with each wave are given in Table 1 with each wave having 2, 4, 15 or 26 nonsynonymous changes. Mutations of the same wave may exhibit nearly identical trajectories (as seen in the overlapped darker line in Fig. 2) yielding a single haplotype. In a slightly different manner, W2 and W4 are represented by multiple groups of mutations of varying frequencies and they all rise and fall in concert.

**Fig. 2.**
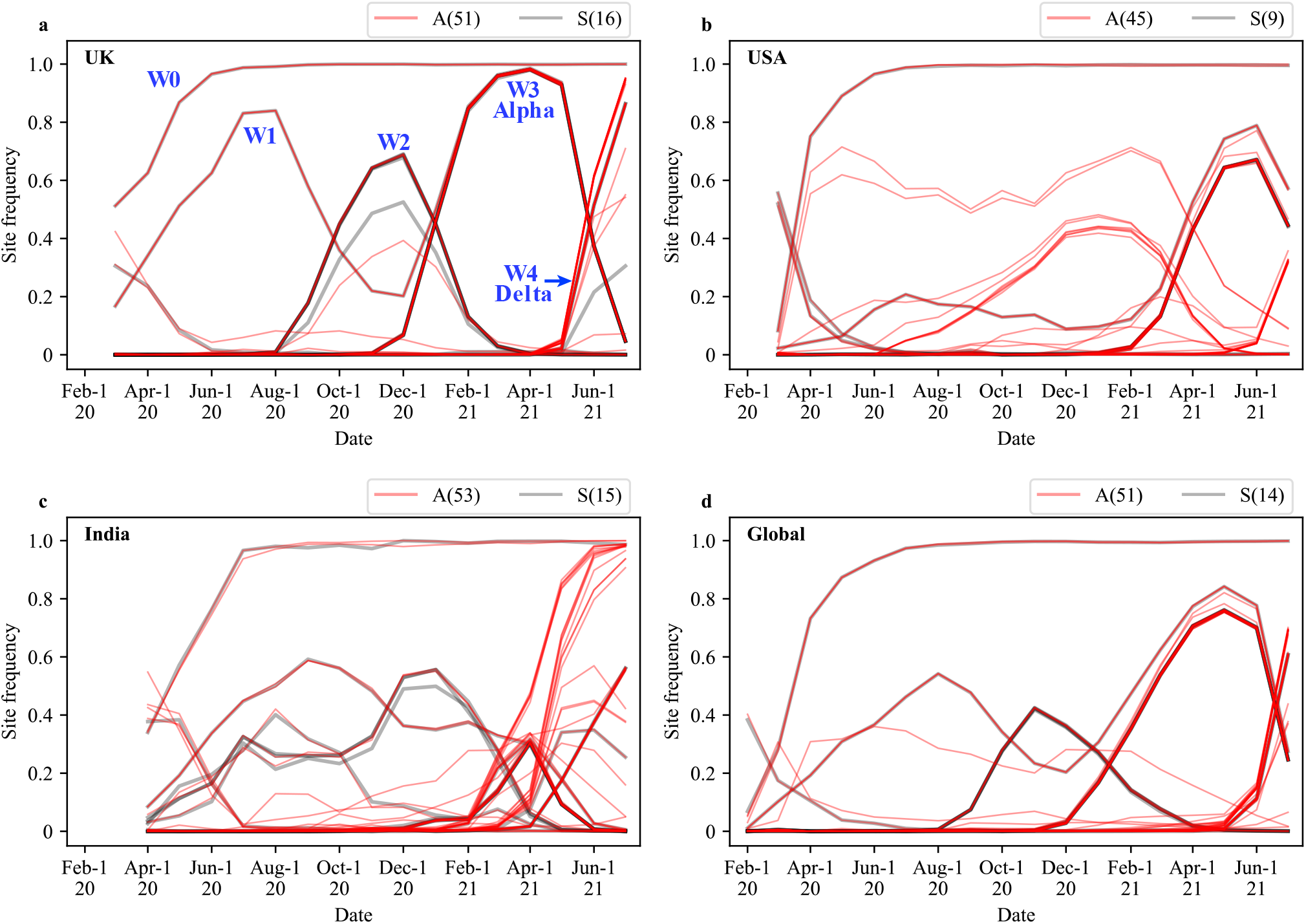
Evolution of SARS-CoV-2 between Feb. 1, 2020 and July 1, 2021 depicted by waves (i.e., successions of “mutation groups”) in UK (**a**), USA (**b**), India (**c**), and Global (**d**). Sequencing data were obtained from the GISAID database. The frequency of variants which reach the frequency cutoff of 0.3 at their peaks are presented. While a curve represents the rise and fall of a variant, each observed curve usually represents multiple curves that overlap completely. In COVID-19, there are 5 waves (W0 to W4). Red lines represent non-synonymous mutations (A), and grey lines are synonymous mutations (S). The number of mutations is shown in the parentheses.

**Table 1.** Number of variant sites associated with each of the five waves in UK.

| Waves | Nonsynonymous (A) | Synonymous (S) | Non-Coding (NC) | A:S:NC |
| --- | --- | --- | --- | --- |
| **W0** | 2 (C14408T, A23403G) | 1 (C3037T) | 1 (C241T) | 2:1:1 |
| **W1** | 2 (G28881A, G28883C) | 1 (G28882A) | 0 | 2:1:0 |
| **W2** | 4 (C21614T, C22227T, C28932T, G29645T) | 5 (T445C, C6286T, G21255C, C26801G, C27944T) | 1 (G204T) | 4:5:1 |
| **W3 (Alpha)** | 15 (see legends) <sup>a</sup> | 5 (C913T, C5986T, C14676T, C15279T, T16176C) | 1 (C27972T) | 15:5:1 |
| **W4 (Delta)** | 26 (see legends) <sup>b</sup> | 3 (C8986T, A11332G, T17040C) | 2 (G210T, G29742T) | 26:3:2 |

<sup>a</sup> C3267T, C5388A, T6954C, A23063T, C23271A, C23604A, C23709T, T24506G, G24914C, G28048T, A28111G, G28280C, A28281T, T28282A, C28977T
<sup>b</sup> G4181T, C6402T, C7124T, C7851T, G9053T, C10029T, A11201G, G15451A, C16466T, C19220T, C21618G, C21846T, G21987A, T22917G, C22995A, C23604G, G24410A, C25469T, T26767C, T27638C, C27752T, C27874T, A28461G, G28881T, G28916T, G29402T

Fig. 3a portrays how mutations appear to evolve in concert even though each emerged sequentially. As shown, the abundance of the four mutations, A-D, differ by at most 11% in the end (5000 to 5555). Therefore, the four curves, if displayed as Fig. 2 does, would overlap almost completely. In the case of W2 and W4, the mutations do differ somewhat in their abundance but all mutations in the same wave still exhibit similar temporal dynamics (see Fig. 2).

**Fig. 3.**
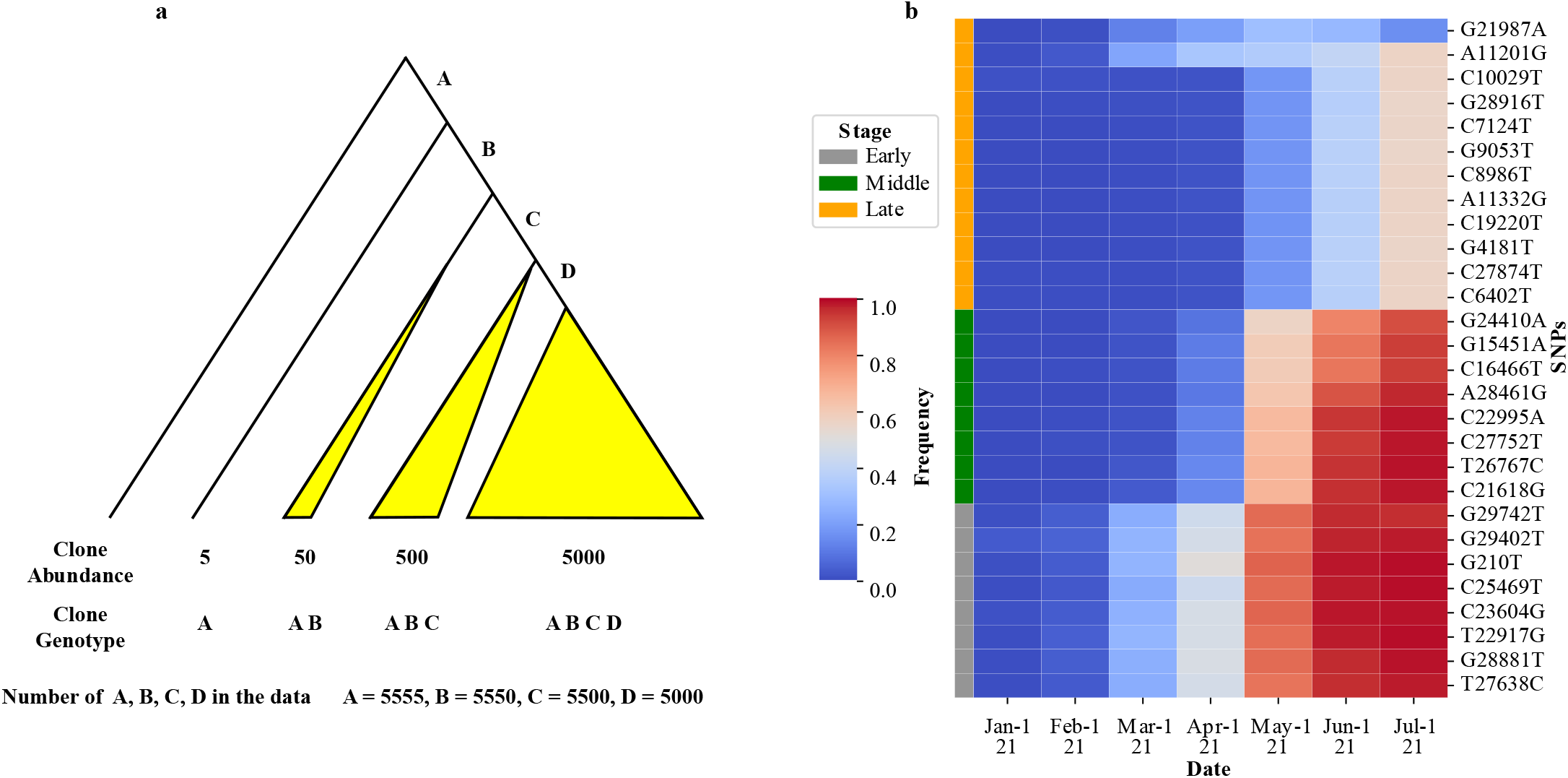
The process of mutation accumulation in clusters. **a**, An illustration of the principle of mutation accumulation. Each of the four mutations, A-D, is acquired step-by-step but a large fitness gain is realized only when all of them are present. As the four mutations would become highly prevalent nearly concurrently, the trajectories of these mutations in Fig. 2 would appear to overlap strongly. **b**, The actual process of mutation accumulation in the evolution of the Delta strain in India. Each row represents a particular nucleotide site and these mutations fall into three groups, labeled E, M and L (early, middle and late, respectively).

The mutations of W0, which include D614G, are fixed by May of 2020 such that subsequent waves are all built on the strain of W0. The dynamics of the 4 W0 mutations (A:S:NC = 2:1:1; see Table 1) illuminates the twin beginnings of COVID-19 as detailed in Ruan et al(Wu 2021). The W0 wave, however, is exceptional. All other strains appear to be driven out by another strain in the next wave. The decline and disappearance of the prevalent strain in later waves can be seen in both the relative and absolute abundance. The continual replacement of one strain by the next one in a series of waves has many implications. An obvious one is that SARS-CoV-2 has become ever more adapted to human conditions, perhaps in response to humans’ social, behavioral and immunological changes.

The most significant observation is the genetic changes of the strains. We note in Table 1 that each new wave is associated with more and more mutations. In the first two waves, there are only two amino acid replacements whereas the number increases to 15 and then 26 replacements in the later Alpha and Delta strains of W3 and W4. In Table 1, the number of amino acid replacements, a proxy of adaptive evolution rate, *R*(*t*), has been increasing as the pandemic progresses. In the remainder of this study, we will focus on the latest Delta strain.

#### 3. The evolution of Delta in UK

W4 of Fig. 2 is the Delta strain. The variants associated with W2 in UK and India are listed in Supplementary Table 5 which reports the monthly trend of variants that exceed 0.3 in their peak frequency. UK has the most extensive sequences and is geographically confined while India is chosen for being the first to report a Delta case(Cherian, et al. 2021; Planas, et al. 2021; Singh, et al. 2021; Zhang, et al. 2021). The Delta strain is defined by the 3 adjacent amino acid (AA) variants in the Spike protein, L452R, T478K, and P681R(Singh, et al. 2021; Tao, et al. 2021). Of course, the Delta strain is far more complicated in its adaptation than the 3 AA’s. The A:S: NC (non-coding) numbers shared between UK and India are 24: 2: 2 while the numbers unique to UK and India are, respectively, 2 : 1: 0 and 4: 1 : 0 (Supplementary Table 5). Delta in UK thus has the 26 : 3 : 2 ratio reported in Table 1. Since the neutral A:S ratio is generally about 2.5 : 1(Li, et al. 1985), there is a large excess of nonsynonymous changes in the Delta strain, indicating its strong adaptive evolution.

By December of 2020, 20 of the 24 AA mutations can be detected in the UK population, albeit in low frequency (≪ 1%). They remain in low frequency through April of 2021 (Fig 2a), during the time the Alpha type was the dominant strain. However, by March, all Delta mutations are seen in UK, suggesting that a complete Delta haplotype has been assembled. Very quickly, the complete haplotype reaches > 1% in May and > 50% by June and > 90% in July in UK (Supplementary Table 5). In short, partial Delta haplotypes spread slowly but exploded out of the gate as soon as the full set of mutations has been in place, as illustrated in Fig. 3a. This pattern would give the impression of simultaneous assembly of all mutations in one strain.

#### 4. The multi-stage evolution of Delta in India

To know how, where and when all the Delta mutations are assembled into the mature product, we analyze the viral sequences from India where the first Delta case was reported(Cherian, et al. 2021; Planas, et al. 2021; Singh, et al. 2021; Tao, et al. 2021). In Supplementary Table 5, the mutations are classified into four groups, E, M, L and R (for early, middle, late and recent) with each mutation’s frequency in each month listed. The pattern in India is graphically presented by the heatmap of Fig. 3b that echoes the hypothetical dynamics of Fig. 3a.

Mutations of each group share the same evolutionary dynamics as illustrated in Fig. 3a-3b. The E, M and L groups are the main variants shared between India and UK, whereas the R groups are very recent mutations found only in India or UK. Among the E, M and L groups, the first detection of variants > 1% happened in February, March and April of 2021, respectively. The first time the variants reach 50% is in April, May and July for the same groups (Fig. 3b). The dynamics of the evolution of Delta in India is portrayed in Fig. 4. Each of the E, M and L groups confers a substantial fitness advantage over the previous group. Crucially, each group has 6 – 10 AA changes. In the M group, for example, the A:S ratio is 8:0.

**Fig. 4.**
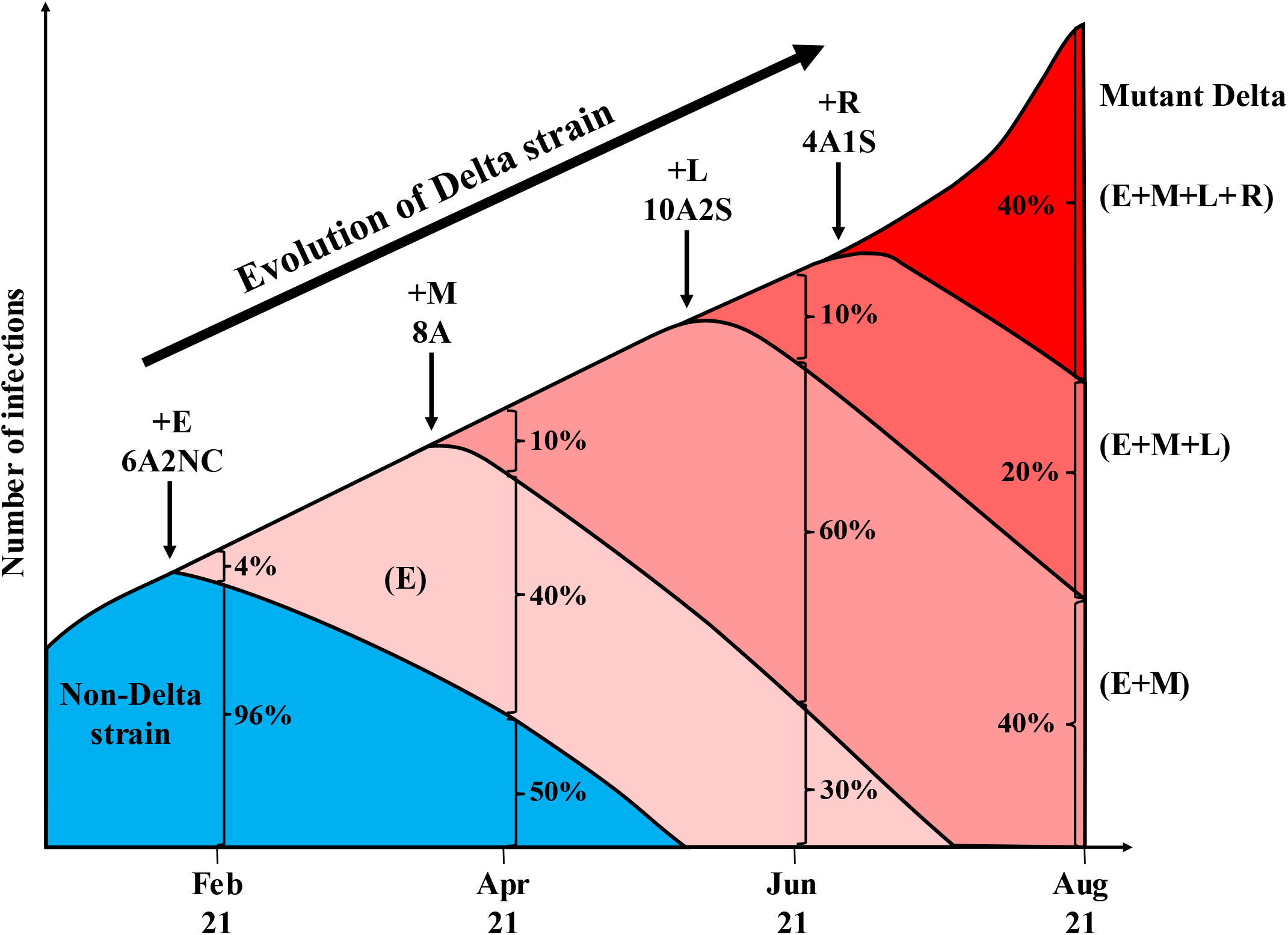
The evolution of the Delta strain in India. The rises (and falls) of five distinct strains are shown in different colors during the evolution of Delta. The light color indicates the non-Delta strains that eventually disappear. The four colors represent the pre-Delta strains (bearing E, E+M and E+M+L mutations) as well as the latest Delta strain bearing E+M+L+R mutations. Note that each of the pre-Delta or Delta strains must start with a single haplotype bearing all the characteristic mutations; hence, the increase in frequency in the beginning must be very substantial. At each time point indicated, the portrayed strains add up to 100%. The size of the entire viral population increases with time but the depiction of the total number corresponds only roughly with the trend. The sets of E, M, L and R mutations are depicted in Fig. 3b shown in the figure as 10A2S for 10 nonsynonymous and 2 synonymous mutations in the group. NC is for non-coding mutations.

The advantage of each group (E, M, L or R) over the previous ones is substantial since the new group must start in one infected person, as shown in Fig. 4. This single infection has to out-compete the earlier haplotype which could account for up to 60% of the total infections at that time. Within the Delta group in India, there have been 4 stages of adaptive shift within a period of 8 months. The A:S ratio suggests that at least 20 of the AA changes confer a fitness advantage and the adaptive mutations have been emerging in quicker successions in February to August of 2021. In every respect, the emergence of the Delta strain is a strong indication that COVID-19 has been experiencing accelerated evolution since early 2021.

#### 5. In the very beginning of the Delta strain – The distribution of mutations among haplotypes

We have so far shown the gradual assembly of the Delta strain in the population. In this last sub-section, we examine the individual haplotypes in the very beginning when even partial Delta haplotypes could not be detected in the population.

According to Supplementary Table 5, the Delta group variants only start to show signs of assembly on Oct. 1, 2020. We now examine the polymorphism data of the 28 sites (E, M, and L group sites) before that day (Fig. 5a). In a sample of 772 sequences collected in India from Sep. 2 to Oct. 1, 2020, 16 of 28 polymorphic sites are singletons (i.e., the variant occurring only once). In most polymorphism data, these rare variants would be scattered among different sequences. In other words, the assembled haplotypes emerge only when the many needed mutations have become common in the population. However, Fig. 5a reveals that a particular haplotype (GISAID: EPI_ISL_2461258) has all of the 16 singleton sites. Furthermore, none of these singletons appear in the 657 sequences collected in the previous month (Supplementary Table 5).

**Fig. 5.**
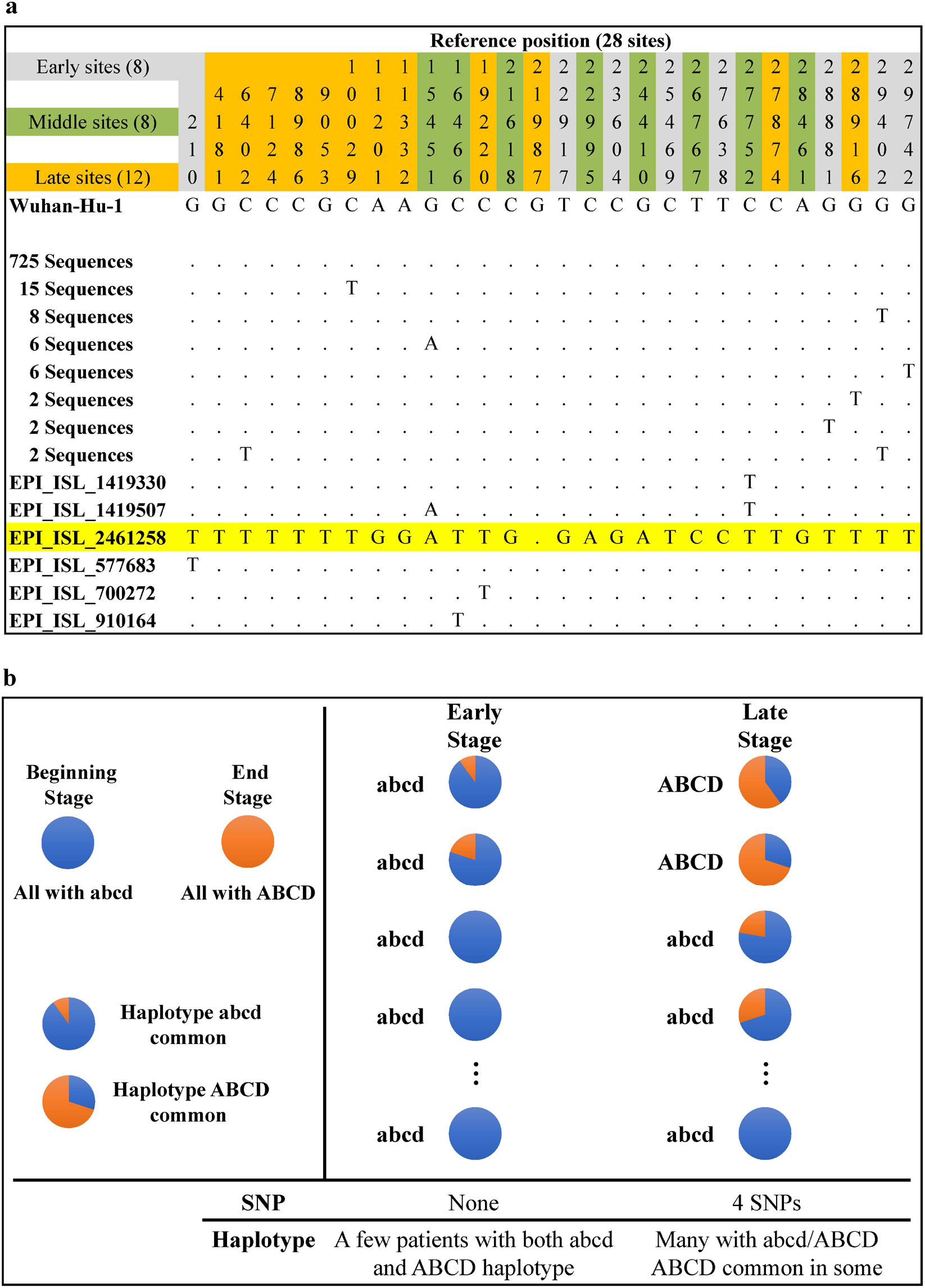
The beginning of haplotype assembly and the distribution of mutations among individuals. The figure attempts to show how the Delta haplotype is first assembled in any individual. **a**, From the data of 2020-9-2 to 2020-10-1, 28 variants are identified from 772 sequences in India. All haplotypes and their occurrences are given. Note that one single haplotype (#2461258) accumulates nearly all the mutants (including most of the singletons at that time) of the Delta strain. **b**, A model on the emergence of a new haplotype (ABCD) from intra-host diversity to become inter-host polymorphism. In this model, the gradual accumulation of mutants happens within hosts, thus creating the impression of sudden appearance of the haplotype (ABCD) between individuals.

Patterns of Fig. 5a raise the following question. How could one haplotype (#2461258) accumulate so many mutants unique to itself? The peculiar phenomenon is plausible if we consider the sequence diversity within each infected individual. The explanation is illustrated in Fig. 5b. The #2461258 haplotype is likely present in many individuals but it is the prevalent haplotype (> 50%) in only one individual among the 772 individuals sampled. In all others, the frequency is < 50% and, hence, is not registered in the database.

Fig. 5a – 5b shows that even the partial assembly of the haplotype #2461258 must have been in progress well before October of 2020. Importantly, the haplotype must first exist as a minor allele within infect hosts when it was being assembled. After Oct. 2020, this haplotype would take several more months to become fully assembled. In short, the Delta strain could be as old as the pandemic itself if we consider the age of the first Delta mutation.

## Discussion

In the mere 20 months, SARS-CoV-2 has been through a rather complex process of evolution with five waves of strains that rise, and often fall subsequently (Fig. 2). The number of mutations in each wave increases from 3-4 in the first two, to 10 and then to 21 and 31 in the last two waves of Alpha and Delta strains (Table 1). Most significant is the A:S ratio that increases from ~1 in the first 3 waves to 3 and then to 8.7 in the last two. In short, SARS-CoV-2 has indeed been experiencing adaptive evolution in an accelerated pace. This accelerated evolution can be even more clearly discerned in the evolution of the latest Delta strain, which proceeds in 4 stages accruing 6 – 12 coding mutations with a high A:S ratio in each stage (Fig. 4). While the assembly of mutations must proceed with one mutation at a time, a large fitness gain is realized only when all mutations are present, as illustrated in Fig. 3.

The main issues are, then, i) how does the virus accumulate these many mutations, many of which conferring a fitness advantage? ii) why does the proportion of advantageous mutations keep increasing? In the theory section of Results, we suggest the runaway evolution via a positive feedback loop. There are indeed a number of forces that can be mutually reinforcing vis-a-vis the process of adaptive evolution. One of them, as Fig. 1 suggests, is a growing *M*(*t*) since small fitness gains are much more likely to spread in large populations (see Eq. 1). The positive feedback loop is expressed as *R*(*t*) → *M*(*t*) and *M*(*t*) → *R*(*t*), which triggers the runaway evolution.

The indication is that the rate of SARS-CoV-2 evolution, *R*(*t*), has been accelerating together with the growth of *M*(*t*). Although *M*(*t*) has been through ups and downs due to human interventions, even highly vaccinated countries have not been able to contain it. We should note that the bidirectional influences of *R*(*t*) → *M*(*t*) and *M*(*t*) → *R*(*t*) can only be realized after a time lag. Hence, an exact correspondence between *R*(*t*) and *M*(*t*) is not expected. By permitting *M*(*t*) to become so large, human societies also permit SARS-CoV-2 to accumulate many mutations. These mutations may have a fitness advantage too small to help when *M*(*t*) is small but the large *M*(*t*) would eventually lead to the emergence of a highly evolved strain like the Delta. In this respect, the global failure in minimizing *M*(*t*) in 2020 may be human societies’ gravest error in dealing with COVID-19.

Finally, there have been intense interests in and out of the academia in unraveling the origin of SARS-CoV-2 as if the origin happened suddenly(Ruan, et al. 2021; Wu, et al. 2021). In contrast, Richard. Dawkins (1986) has famously promoted the Blind Watchmaker argument for the evolution of complex biological traits through a long process of step-by-step evolution(Dawkins 1986). By the same argument, it has been suggested that SARS-CoV-2 must have been through a long process of stepwise adaptive evolution(Wu, et al. 2021). This process could take place only in a natural setting, the main features of which having been outline in a previous model(Ruan, et al. 2021). In this study, we present the empirical evidence for the complex evolutionary process of SARS-CoV-2 in 2020-2021. Even between two strains of the same human host (Alpha vs. Delta), there have been 41 amino acid changes. It is hence reasonable to suggest that the host shift from animals to humans likely require even more changes, which can only happen step by step in nature.

## Materials and Methods

### Data collection and pre-processing

We download SARS-CoV-2 genomes from the GISAID database(Elbe and Buckland-Merrett 2017) as of July 5, 2021 with the following download options: (1) “complete”: genomes > 29,000nt; (2) “low coverage excel”: exclude viruses with >5% Ns (undefined base). All animal isolate strains were removed and 2,063,459 SARS-CoV-2 genomes were retained. In addition, we filtered out sequences without collection date or with an implausible collection date and retained 1,853,355 genomes for downstream analysis.

### Sequence alignment, SNP calling and annotation

We aligned these 1,853,355 genome sequences to the reference sequence (Wuhan-Hu-1(Wu, et al. 2020), GenBank: NC_045512, GISAID: EPI_ISL_402125) using MAFFT (--auto -- keeplength)(Katoh and Standley 2013). We used snp-sites (-v)(Page, et al. 2016) to identify SNPs and BCFtools (merge --force-samples -O v)(Li, et al. 2009) to merge the vcf files. Interestingly, although the reference genome size is only 29,903 nt, we found 67,650 SNP sites from these 1,853,355 genome sequences, indicating multiple substitutions at the same sites. We annotated the 67,650 SNP sites by ANNOVAR(Yang and Wang 2015).

### Analysis of SNPs

In this study, we track the variant frequency at each variable site (e.g., C→T). If we set the cutoff by ignoring variants that fail to reach 0.3 in frequency, there are usually fewer than 100 variants to keep track of. To observe the changes in site frequencies in a geographic region, we group the data into bins each covering a one-month period. Only variants reaching the frequency cutoff of 0.3 at their peaks are retained. For the retained variants, we grouped them by pairwise Pearson coefficients with the cutoff of 0.9 (i.e., if the Pearson coefficient across the time span between two variants is equal or greater than 0.9, they would belong in the same group). With this cutoff, there are 5 major waves (Fig. 2).

## Data availability

All genomic data in this study are available, on registration, from GISAID (https://www.gisaid.org/).

## Acknowledgments

We thank all those who have contributed sequences to the GISAID (Global Initiative on Sharing All Influenza Data) database (https://www.gisaid.org/).

## References

Andersen KG, Rambaut A, Lipkin WI, Holmes EC, Garry RF. 2020. The proximal origin of SARS-CoV-2. Nature Medicine 26:450–452.

Cherian S, Potdar V, Jadhav S, Yadav P, Gupta N, Das M, Rakshit P, Singh S, Abraham P, Panda S. 2021. Convergent evolution of SARS-CoV-2 spike mutations, L452R, E484Q and P681R, in the second wave of COVID-19 in Maharashtra, India. bioRxiv:2021.2004.2022.440932.

Choi B, Choudhary MC, Regan J, Sparks JA, Padera RF, Qiu X, Solomon IH, Kuo HH, Boucau J, Bowman K, et al. 2020. Persistence and Evolution of SARS-CoV-2 in an Immunocompromised Host. N Engl J Med 383:2291–2293.

Crow JF, Kimura M. 1970. An Introduction to Population Genetics Theory: Harper & Row.

Cui J, Li F, Shi ZL. 2019. Origin and evolution of pathogenic coronaviruses. Nature Reviews: Microbiology 17:181–192.

Dawkins R. 1986. The Blind Watchmaker: Longman Scientific & Technical.

Dejnirattisai W, Zhou D, Supasa P, Liu C, Mentzer AJ, Ginn HM, Zhao Y, Duyvesteyn HME, Tuekprakhon A, Nutalai R, et al. 2021. Antibody evasion by the P.1 strain of SARS-CoV-2. Cell 184:2939–2954 e2939.

Deng X, Garcia-Knight MA, Khalid MM, Servellita V, Wang C, Morris MK, Sotomayor-Gonzalez A, Glasner DR, Reyes KR, Gliwa AS, et al. 2021. Transmission, infectivity, and neutralization of a spike L452R SARS-CoV-2 variant. Cell:S0092-8674(0021)00505-00505.

Elbe S, Buckland-Merrett G. 2017. Data, disease and diplomacy: GISAID’s innovative contribution to global health. Glob Chall 1:33–46.

Eyre-Walker A. 2006. The genomic rate of adaptive evolution. Trends in Ecology & Evolution 21:569–575.

Forster P, Forster L, Renfrew C, Forster M. 2020. Phylogenetic network analysis of SARS-CoV-2 genomes. Proc Natl Acad Sci U S A 117:9241–9243.

Guo H, Hu BJ, Yang XL, Zeng LP, Li B, Ouyang S, Shi ZL. 2020. Evolutionary Arms Race between Virus and Host Drives Genetic Diversity in Bat Severe Acute Respiratory Syndrome-Related Coronavirus Spike Genes. Journal of Virology 94:e00902–00920.

Hou YJ, Chiba S, Halfmann P, Ehre C, Kuroda M, Dinnon KH, 3rd, Leist SR, Schafer A, Nakajima N, Takahashi K, et al. 2020. SARS-CoV-2 D614G variant exhibits efficient replication ex vivo and transmission in vivo. Science 370:1464–1468.

Kajan GL, Doszpoly A, Tarjan ZL, Vidovszky MZ, Papp T. 2020. Virus-Host Coevolution with a Focus on Animal and Human DNA Viruses. Journal of Molecular Evolution 88:41–56.

Katoh K, Standley DM. 2013. MAFFT multiple sequence alignment software version 7: improvements in performance and usability. Molecular Biology and Evolution 30:772–780.

Kemp SA, Collier DA, Datir RP, Ferreira I, Gayed S, Jahun A, Hosmillo M, Rees-Spear C, Mlcochova P, Lumb IU, et al. 2021. SARS-CoV-2 evolution during treatment of chronic infection. Nature 592:277–282.

Kimura M. 1969. The number of heterozygous nucleotide sites maintained in a finite population due to steady flux of mutations. Genetics 61:893–903.

Kimura M, Crow JF. 1964. The Number of Alleles That Can Be Maintained in a Finite Population. Genetics 49:725–738.

Korber B, Fischer WM, Gnanakaran S, Yoon H, Theiler J, Abfalterer W, Hengartner N, Giorgi EE, Bhattacharya T, Foley B, et al. 2020. Tracking Changes in SARS-CoV-2 Spike: Evidence that D614G Increases Infectivity of the COVID-19 Virus. Cell 182:812–827 e819.

Kumar S, Tao Q, Weaver S, Sanderford M, Caraballo-Ortiz MA, Sharma S, Pond SLK, Miura S. 2021. An Evolutionary Portrait of the Progenitor SARS-CoV-2 and Its Dominant Offshoots in COVID-19 Pandemic. Molecular Biology and Evolution 38:3046–3059.

Li H, Handsaker B, Wysoker A, Fennell T, Ruan J, Homer N, Marth G, Abecasis G, Durbin R, Genome Project Data Processing S. 2009. The Sequence Alignment/Map format and SAMtools. Bioinformatics 25:2078–2079.

Li Q, Wu J, Nie J, Zhang L, Hao H, Liu S, Zhao C, Zhang Q, Liu H, Nie L, et al. 2020. The Impact of Mutations in SARS-CoV-2 Spike on Viral Infectivity and Antigenicity. Cell 182:1284–1294 e1289.

Li WH, Wu CI, Luo CC. 1985. A new method for estimating synonymous and nonsynonymous rates of nucleotide substitution considering the relative likelihood of nucleotide and codon changes. Molecular Biology and Evolution 2:150–174.

Lytras S, Xia W, Hughes J, Jiang X, Robertson DL. 2021. The animal origin of SARS-CoV-2. Science 373:968–970.

Page AJ, Taylor B, Delaney AJ, Soares J, Seemann T, Keane JA, Harris SR. 2016. SNP-sites: rapid efficient extraction of SNPs from multi-FASTA alignments. Microb Genom 2:e000056.

Parrish CR, Holmes EC, Morens DM, Park EC, Burke DS, Calisher CH, Laughlin CA, Saif LJ, Daszak P. 2008. Cross-species virus transmission and the emergence of new epidemic diseases. Microbiology and Molecular Biology Reviews 72:457–470.

Peschel A, Sahl HG. 2006. The co-evolution of host cationic antimicrobial peptides and microbial resistance. Nature Reviews: Microbiology 4:529–536.

Planas D, Veyer D, Baidaliuk A, Staropoli I, Guivel-Benhassine F, Rajah MM, Planchais C, Porrot F, Robillard N, Puech J, et al. 2021. Reduced sensitivity of SARS-CoV-2 variant Delta to antibody neutralization. Nature 596:276–280.

Plante JA, Liu Y, Liu J, Xia H, Johnson BA, Lokugamage KG, Zhang X, Muruato AE, Zou J, Fontes-Garfias CR, et al. 2021. Spike mutation D614G alters SARS-CoV-2 fitness. Nature 592:116–121.

Plowright RK, Parrish CR, McCallum H, Hudson PJ, Ko AI, Graham AL, Lloyd-Smith JO. 2017. Pathways to zoonotic spillover. Nature Reviews: Microbiology 15:502–510.

Rambaut A, Holmes EC, O’Toole A, Hill V, McCrone JT, Ruis C, du Plessis L, Pybus OG. 2020. A dynamic nomenclature proposal for SARS-CoV-2 lineages to assist genomic epidemiology. Nat Microbiol 5:1403–1407.

Ruan Y, Wang H, Chen B, Wen H, Wu CI. 2020. Mutations Beget More Mutations-Rapid Evolution of Mutation Rate in Response to the Risk of Runaway Accumulation. Molecular Biology and Evolution 37:1007–1019.

Ruan Y, Wang H, Zhang L, Wen H, Wu C-I. 2020. Sex, fitness decline and recombination – Muller’s ratchet vs. Ohta’s ratchet. bioRxiv:2020.2008.2006.240713.

Ruan Y, Wen H, He X, Wu CI. 2021. A theoretical exploration of the origin and early evolution of a pandemic. Sci Bull (Beijing) 66:1022–1029.

Sashittal P, Luo Y, Peng J, El-Kebir M. 2020. Characterization of SARS-CoV-2 viral diversity within and across hosts. bioRxiv:2020.2005.2007.083410.

Singh J, Rahman SA, Ehtesham NZ, Hira S, Hasnain SE. 2021. SARS-CoV-2 variants of concern are emerging in India. Nature Medicine 27:1131–1133.

Tang X, Wu C, Li X, Song Y, Yao X, Wu X, Duan Y, Zhang H, Wang Y, Qian Z, et al. 2020a. On the origin and continuing evolution of SARS-CoV-2. National Science Review 7:1012–1023.

Tang X, Wu C, Li X, Song Y, Yao X, Wu X, Duan Y, Zhang H, Wang Y, Qian Z, et al. 2020b. On the origin and continuing evolution of SARS-CoV-2. National Science Review 7:1012–1023.

Tang X, Ying R, Yao X, Li G, Wu C, Tang Y, Li Z, Kuang B, Wu F, Chi C, et al. 2021. Evolutionary analysis and lineage designation of SARS-CoV-2 genomes. Sci Bull (Beijing) 66:2297–2311.

Tao K, Tzou PL, Nouhin J, Gupta RK, de Oliveira T, Kosakovsky Pond SL, Fera D, Shafer RW. 2021. The biological and clinical significance of emerging SARS-CoV-2 variants. Nature Reviews: Genetics 22:757–773.

Volz E, Hill V, McCrone JT, Price A, Jorgensen D, O’Toole A, Southgate J, Johnson R, Jackson B, Nascimento FF, et al. 2021. Evaluating the Effects of SARS-CoV-2 Spike Mutation D614G on Transmissibility and Pathogenicity. Cell 184:64–75 e11.

Wu C-I. 2021. The twin-beginnings of COVID-19 in Asia and Europe – One prevails quickly. Research Square.

Wu CI, Wen H, Lu J, Su XD, Hughes AC, Zhai W, Chen C, Chen H, Li M, Song S, et al. 2021. On the origin of SARS-CoV-2-The blind watchmaker argument. Sci China Life Sci 64:1560–1563.

Wu F, Zhao S, Yu B, Chen YM, Wang W, Song ZG, Hu Y, Tao ZW, Tian JH, Pei YY, et al. 2020. A new coronavirus associated with human respiratory disease in China. Nature 579:265–269.

Yang H, Wang K. 2015. Genomic variant annotation and prioritization with ANNOVAR and wANNOVAR. Nat Protoc 10:1556–1566.

Zhang J, Xiao T, Cai Y, Lavine CL, Peng H, Zhu H, Anand K, Tong P, Gautam A, Mayer ML, et al. 2021. Membrane fusion and immune evasion by the spike protein of SARS-CoV-2 Delta variant. Science:eabl9463.

